# BCG vaccination of Diversity Outbred mice induces cross-reactive antibodies to SARS-CoV-2 spike protein

**DOI:** 10.1101/2022.04.18.488640

**Authors:** Aubrey G. Specht, Sherry L. Kurtz, Karen L. Elkins, Harrison Specht, Gillian Beamer

## Abstract

The Bacillus Calmette-Guérin (BCG) vaccine, the only vaccine against tuberculosis, induces cross-protection against pathogens unrelated to *Mycobacterium*, including viruses. Epidemiological studies have identified potential benefits of BCG vaccination against SARS-CoV-2 infection. While BCG’s heterologous effects have been widely attributable to *trained immunity*, we hypothesized BCG vaccination could induce cross-reactive antibodies against the spike protein of SARS-CoV-2 Wuhan-Hu-1. The concentration of IgG reactive to SARS-CoV-2 spike protein from the sera of BCG-vaccinated, Diversity Outbred (DO) mice and C57BL/6J inbred mice was measured using ELISA. Sera from 10/15 BCG-vaccinated DO mice possessed more IgG reactive to recombinant spike protein than sera from BCG-vaccinated C57BL/6J mice and unvaccinated DO mice. Amino acid sequences common to BCG cell wall/membrane proteins and SARS-CoV-2 spike protein were identified as potential antigen candidates for future study. These results imply a humoral mechanism, influenced by genotype, by which BCG vaccination could confer immunity to SARS-CoV-2.

**Graphic Abstract:** 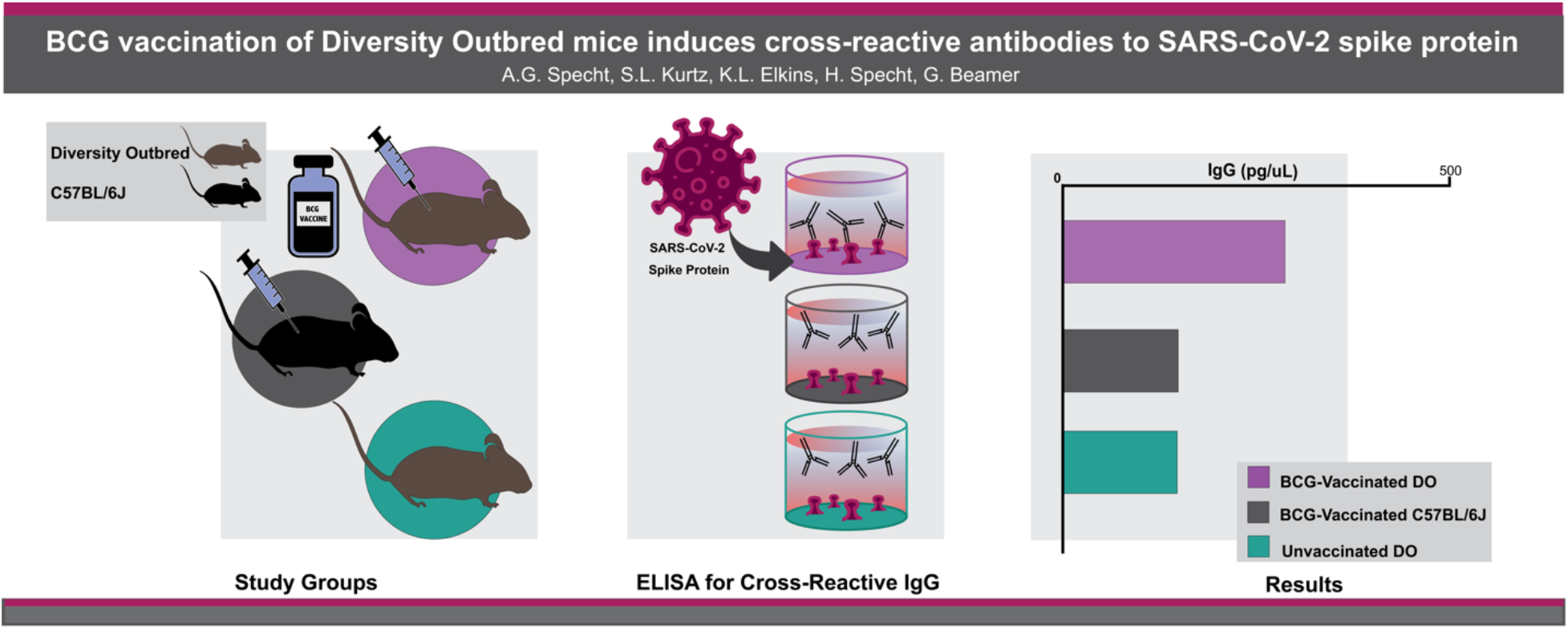

## Introduction

### Cross-Protective Benefits of the BCG vaccine

Bacillus Calmette-Guérin (BCG) is a live attenuated strain of *Mycobacterium bovis* and used to vaccinate 130 million children in 154 countries each year against tuberculosis (TB) [1]. However, BCG’s impact extends beyond TB. Starting in 1927, epidemiological and basic research indicates that immunity induced by BCG confers cross-protection against other pathogens, including viruses [2,3].

### Epidemiological, Clinical, and Mechanistic Evidence for BCG’s Cross-Protection Against SARS-CoV-2

The SARS-CoV-2 virus did not affect countries equally. During the early months of the pandemic, epidemiological analyses showed that a 10% increase in BCG index, an estimate for a country’s BCG vaccination use, was associated with a 10.4% decrease in COVID-19 mortality [4]. This intriguing finding led to clinical trials, seeking prospective validation. In the phase III trial ACTIVATE (NCT03296423), BCG revaccination of elderly patients (> 65 years) yielded 68% risk reduction for total COVID-19 clinical and microbiological diagnoses in the 6 months following vaccination [5]. On-going phase III clinical trials, such as BRACE (NCT04327206) and BATTLE (NCT04369794), are assessing BCG’s protection against SARS-CoV-2 infection and its use as a therapeutic tool in patients currently infected with SARS-CoV-2, respectively [6]. Proposed mechanisms for BCGs cross-protection include: 1) trained immunity of the innate immune system, driven by monocytes and natural killer (NK) cells; 2) T cell lymphocyte-dependent activation of innate immunity and adaptive immunity; and 3) antibody cross-reactivity to a wide array of antigens [1].

### Humoral Immunity’s Role in Preventing Infection

To infect host cells, SARS-CoV-2’s surface spike glycoproteins bind the angiotensin-converting enzyme 2 (ACE-2) receptors on host cells [7]. Neutralizing antibodies prevent this initial binding [8], demonstrating a role for B-cell derived (humoral) immunity against SARS-CoV-2.

The role B cells and antibodies play in the cross-protective benefits of BCG against SARS-CoV-2 is currently debated. *In silico* analysis has found 690 BCG B-cell epitopes similar to SARS-CoV-2 B-cell epitopes, including epitopes in the RBD of the spike protein [9]. Using immunohistochemistry, another study found that an antibody directed against the SARS-CoV-2 envelope protein also bound the LytR C protein found in *M. bovis* [10]. However, results from an *in vivo* study in the BALB/c inbred mouse strain appear contradictory, as BCG vaccination did not produce cross-reactive neutralizing antibodies against SARS-CoV-2 [11].

Here, we investigated BCG-induced antibodies against SARS-CoV-2 spike glycoprotein. *In silico* analysis showed homology between predicted antibody epitopes on the SARS-CoV-2 Spike protein and the proteome of BCG. Further, we measured cross-reactive antibodies to the SARS-CoV-2 Spike protein in different mouse strains (*i.e*., Diversity Outbred (DO) mice and C57BL/6J inbred mice). We compared the level of SARS-CoV-2 reactive antibodies in the sera of BCG-vaccinated, outbred Diversity Outbred (DO) and inbred C57BL/6J mice with the sera of unvaccinated DO mice.

## Materials and Methods

### *In Silico* Analysis of Homology

Bepipred Linear Epitope Prediction 2.0 from IEBD was used to select predicted antibody binding epitopes (>5 amino acids long) on the SARS-CoV-2 spike protein (SwissProt: P0DTC2). Basic Local Alignment Search Tool (BLAST) from NCBI was used to align the selected epitopes with the Swiss-Prot reviewed proteome of BCG strain Pasteur (*Mycobacterium bovis* (strain BCG / Pasteur 1173P2). The results were restricted to only matches with >70% ID and alignment length ≥7 amino acids, based on previous research on homologous peptide length and T cell activation [12]. Pentamers with 100% ID and 6-mer with 83.3% ID were also included, because antibody epitopes can range from 4-12 amino acids [13]. The BCG protein and cellular component, as described on the UniprotKB database, was obtained for each peptide fitting the above criteria. Figures 4a and 4b were created using the R software package *ggplot2*.

### BCG-vaccinated mice

Studies were performed under protocols approved by the Animal Care and Use Committee (ACUC) of CBER/FDA protocol #2011-14. Briefly, female C57BL/6J and female Diversity Outbred (DO) mice, purchased from Jackson Laboratories (Bar Harbor, ME), were used at 6-10 weeks of age. All mice were housed in groups by strain in microisolator cages and given autoclaved food and water *ad libitum*. A single, frozen vial of BCG was thawed and diluted in sterile PBS. Mice were vaccinated intradermally or intravenously with 10^5^ BCG Pasteur colony forming units (Table 1). Eight weeks later, mice were euthanized and blood was obtained via cardiac puncture with a heparinized 1 ml syringe and 26g needle. Sera were separated using Sarstedt microtubes (Fisher Scientific, Pittsburg, PA) [14], stored at −80°C and shipped from Food and Drug Administration (Silver Spring, MD) to Cummings School of Veterinary Medicine (Grafton, MA) for assays.

**Table 1.** Test Sera: Mouse identification and BCG vaccination route.

| Group ID # | Mouse Strain | Vaccine | Vaccine Administration Route | Mouse # | Age (months) |
| --- | --- | --- | --- | --- | --- |
| 17.3 | C57BL/6J | BCG | Intradermal | M1-5 | 3.5 – 4.5 |
| 17.5 | C57BL/6J | BCG | Intravenous | M1-5 | 3.5 – 4.5 |
| 17.11 | DO | BCG | Intradermal | M1-5 | 3.5 – 4.5 |
| 17.15 | DO | BCG | Intravenous | M1-5 | 3.5 – 4.5 |
| 17.16 | DO | BCG | Intravenous | M1-5 | 3.5 – 4.5 |
| GB07.7 | DO | Unvaccinated | n/a | #933 | 5 |
| GB07.6 | DO | Unvaccinated | n/a | #730<br>#731<br>#732<br>#734 | 18 |

### Unvaccinated mice

All procedures on live mice were approved by Tufts University’s Institutional Animal Care and Use Committee (IACUC) protocols G2015-33 or G2018-02, or the CEBR/FDA protocol #2011-14. Female DO mice, purchased from Jackson Laboratories, were housed socially, and provided sterile food and water *ad libitum*. Unvaccinated mice received no treatment. Mice were euthanized, blood was collected via cardiac puncture, allowed to clot, centrifuged and serum collected, and stored at −80°C until use (Table 1).

### In-house ELISA to SARS-CoV-2 Antigens

We optimized an in-house ELISA to detect cross-reactive IgG antibodies in mouse sera. On day 1, Costar 96-well plates were coated in 1 μg/mL recombinant SARS-CoV-2 Spike protein (“rSpike,” BEI Resources NR-52308) diluted in PBS. Half of the plate was left uncoated to measure nonspecific interactions [15]. Plates were incubated overnight at 4°C. On day 2, plates were blocked with 1% Oxoid skim milk + Sodium Azide for 2 hours at 25 4°C and then washed six times with 1X PBS-Tween. To create the standard curve, BEI Resources (NR-616) monoclonal anti-SARS-CoV spike protein was serially diluted starting at 10,000 pg/mL in 1% BSA (Supplemental Figure 1). As assay positive control, aliquots of 240C monoclonal antibody were prepared in 1% BSA at 20,000 pg/mL and frozen at −80°C for consistency across ELISAs (Supplemental Figure 1). All mouse sera were initially diluted 1:500 in 1% BSA and tested in triplicate. Negative control wells contained 1% BSA. Plates were incubated overnight at 44°C.

On day 3, plates were washed six times with 1X PBS-Tween, and horseradish peroxidase (HRP) conjugated goat anti-mouse IgG (Rockland #610-13) was used to detect SARS-CoV-2-bound IgG. Plates were washed six times with 1X PBS-Tween. Plates were incubated 20 minutes with 3,3′,5,5′-Tetramethylbenzidine (TMB), and the colorimetric reaction was stopped with 0.25M HCl. Absorbance was read at 450 nm using a BioTek plate reader. Concentration of bound-IgG was computed based on a 4-parameter logistic (4PL) regression model of the standard curve using the program Gen5.

#### Assay control

On each plate, an independent assay control was run to assess the quality of the detection antibody and colorimetric reaction, as follows: On day 1, the well was coated with 5 μg/mL of a mixture containing recombinant ESAT-6 and CFP-10 (BEI Resources NR-49424, NR-49425). On day 2, sera from CFP-10-vaccinated mice, diluted 1:100 in 1% BSA, were incubated in the well. On day 3, the same detection antibody, goat anti-mouse IgG-HRP, was added followed by the addition of TMB and then HCl, as described above.

#### Statistical Analysis

Graphs were made using Prism GraphPad version 9.1.0. Standard curve limit of detection was calculated based on the sum of the mean blank optical density and three times the standard deviation based on 6 blanks. Histograms and Student’s *t*-tests were performed using R.

## Results/Discussion

Early in the COVID-19 pandemic, epidemiological evidence suggested cross-protection between the BCG vaccine and SARS-CoV-2 infection, fueling investigation into possible mechanisms such as cross-reactive antibodies.

### *In Silico* Analysis of Homology

Potential cross-reactive antibody binding sites were predicted by *in silico* comparison of antibody epitopes on the SARS-CoV-2 Spike protein and the proteome of BCG. Unlike other *in silico* analysis, only predicted linear epitopes >5 amino acids from the RBD were searched against a database of Swiss-Prot reviewed BCG Pasteur proteins. The resulting homologous BCG proteins were characterized by their location on/in *M. bovis*. Bepipred Linear Epitope Prediction 2.0 from IEBD found 25 potential epitopes ≥5 amino acids long on the SARS-CoV-2 spike protein. A BLAST search comparing epitopes with the BCG strain Pasteur proteome yielded 186 peptides (percent ID ≥70% and alignment length ≥7 amino acids, percent ID ≥83.3% and 6-mer, percent ID 100% and 5-mer). The frequency of each percent ID, alignment length pair is described in Figure 1a. Of note, one heptamer had 100% identity with a BCG protein. As only Swiss-Prot reviewed proteins from the BCG Pasteur proteome were used in the BLAST search, all 186 peptides matched with a described BCG protein. Of those BCG proteins, the cellular component was known for 59.7%. Most BCG proteins were cytoplasmic (Figure 1b); however, five proteins are known to be located on the cell surface (Figure 1c). These analyses imply shared sequences of amino acids between BCG surface proteins and the SARS-CoV-2 spike protein.

**Figure 1.**
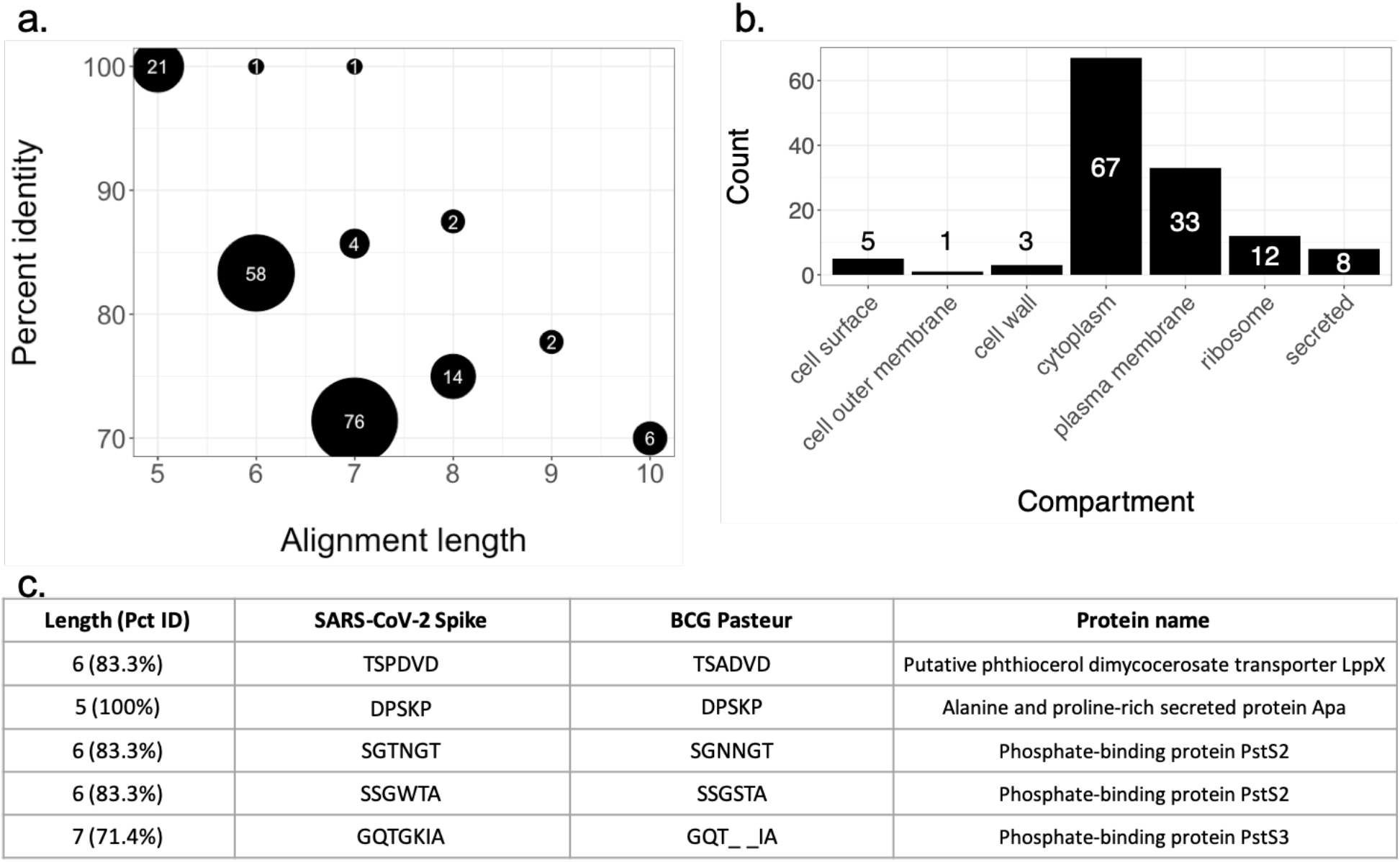
*In silico* analysis of SARS-CoV-2 spike protein epitopes homologous to BCG proteins. 1a. Frequency of alignment length and percent identity pair. Peptides with homology to BCG proteins were restricted to those ≥70% identity and ≥7 amino acids. Hexamers with ≥83% identity and pentamers with 100% identity were also included. Frequency of each pair (alignment length, percent identity) is denoted by size of the data point. 1b. Known cellular components of BCG proteins with SARS-CoV-2 homology. Peptides fitting the above described criteria were further matched with name of the corresponding Swiss Prot *Mycobacterium bovis* strain BCG Pasteur protein. Those proteins’ cellular components were described, if known. The frequency of the cellular components was described. 1c. Homologous peptides restricted to BCG proteins located on the cell surface. The five BCG Pasteur proteins known to be located on the cell surface from 1b were identified and their sequences compared with the SARS-CoV-2 Spike protein.

### Concentration of Cross-reactive IgG to SARS-CoV-2 Antigens using ELISA

The concentration of IgG binding SARS-CoV-2 rSpike protein was compared between Diversity Outbred mice vaccinated with BCG, C57BL/6J mice vaccinated with BCG, and unvaccinated Diversity Outbred mice (Figure 2). Sera from BCG-vaccinated C57BL/6J mice (n=10) were compared with sera from BCG-vaccinated DO mice (n=15) and unvaccinated DO mice (n=5). The signal from uncoated wells incubated with the sera was undetectable. As the unvaccinated DO mice have not been challenged by any pathogen, the signal and resulting concentration of IgG produced by the ELISA is a presumed noise, due to non-specific interactions between normal serum components with assay reagents.

**Figure 2:**
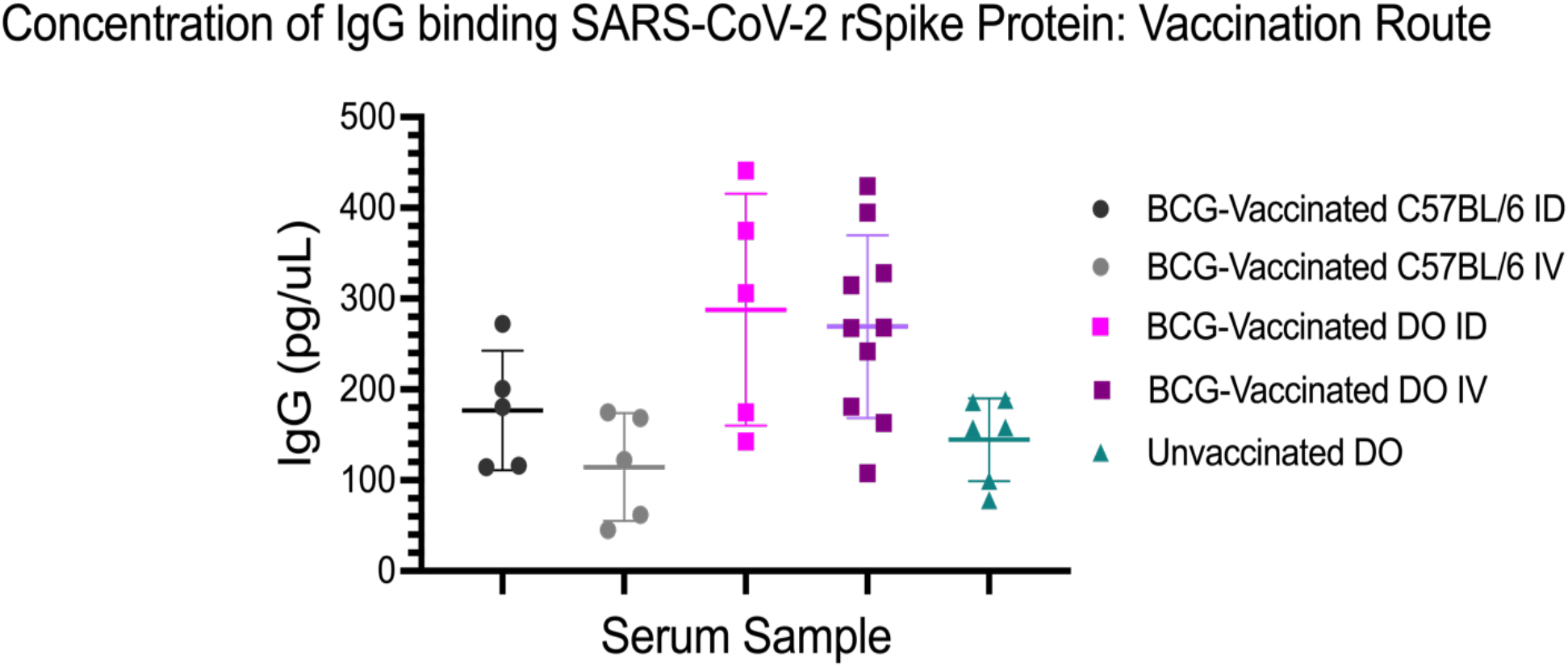
Concentration of IgG binding SARS-CoV-2 rSpike protein by vaccination route and mouse strain. Each data point is the mean concentration based on technical triplicates at 1:500 dilution. C57BL/6J mice vaccinated intradermally (ID) (n=5) and intravenously (IV) (n=5) were compared with DO (J:DO) mice vaccinated ID (n=5), IV (n=10) and unvaccinated (n=5). The Student’s *t*-test was used to determine if the means of two groups are significantly different. IgG concentrations for BCG-vaccinated DO mouse sera (both ID and IV) were significantly greater than both BCG-vaccinated C57BL/6J (both ID and IV) and unvaccinated DO mouse sera.

A subset of the BCG-vaccinated DO mice (10/15 or 67%) produced more antibodies reactive to the recombinant spike protein than unvaccinated DO mice and BCG-vaccinated C57BL/6J mice. The mean IgG concentration reactive to the spike protein in BCG-vaccinated DO mice (292 pg/mL) was significantly different from the mean IgG concentration in unvaccinated DO mice (146 pg/mL; Student’s *t*-test, p,0.001) and in BCG-vaccinated C57BL/6J mice (144 pg/mL, p,0.001). The IgG concentration in BCG-vaccinated C57BL/6J was not significantly different from unvaccinated DO mice (p=0.9171). This interesting difference between BCG-vaccinated DO mice and BCG-vaccinated C57BL/6J mice could reflect assay sensitivity, reagents that better detect antibodies in DO mice, or potentially genetic differences that control humoral immune responses.

The results were examined by vaccine route (Figure 2). Sera from C57BL/6J mice vaccinated intradermally (ID) (n=5) and intravenously (IV) (n=5) were compared with sera from DO mice vaccinated ID (n=5), IV (n=10) and unvaccinated mice (n=5). Neither the difference in IgG concentration between C57BL/6J mice vaccinated IV (177 pg/mL) and ID (115 pg/mL) nor the difference between DO mice vaccinated IV (288 pg/mL) and ID (269 pg/mL), was significant (p=0.1542, p=0.7607). The route of vaccination did not impact concentration of IgG reactive to the recombinant spike protein for either DO or C57BL/6J mice.

Based on the differences observed, we hypothesize that genetic background contributes to the ability of 67% of DO mice to generate BCG-induced IgG cross-reactive to SARS-CoV-2 spike protein. DO mice are as genetically diverse as the human population and may better model the range of response to BCG vaccination [16]. Differences in inbred mouse strains used by researchers may account for contradictory results from studies assessing SARS-CoV-2 reactive antibody levels. For example, Kandeil *et al*. found BCG vaccination did *not* produce detectable cross-reactive neutralizing antibodies against SARS-CoV-2 using the inbred BALB/c strain [11].

Future research could explore this study’s limitations by characterizing the binding site of the cross-reactive IgG antibodies and assessing the antibodies for neutralizing efficacy, or by investigating the genetic basis for the cross-reactive humoral response in DO mice. To the first point, affinity purification mass spectrometry (AP-MS) could confirm antibody concentration and purify the cross-reactive antibodies for affinity testing against peptides identified by our *in silico* analysis. To the second point, genotype-phenotype association studies, such as quantitative trait locus mapping, could be performed on larger sample sizes to identify gene candidates and polymorphisms.

## Conclusion

Using an ELISA, we found that a subgroup of BCG-vaccinated Diversity Outbred mice produced a concentration of cross-reactive IgG antibodies to the spike protein on SARS-CoV-2 above that of unvaccinated mice, and interestingly BCG-vaccinated inbred C57BL/6J mice.

## Abbreviations

(DO): Diversity Outbred

## Acknowledgements

Melanie Ginese

The authors declare that they have no known competing financial interests or personal relationships that could have appeared to influence the work reported in this paper.

Experimental Design: A.S., G.B.
Sample collection: K.E., S.K., G.B.
ELISAs: A.S., G.B.
*In silico* analysis: A.S., H.S.
Data analysis: A.S., H.S.
Writing and editing: A.S., G.B., H.S., K.E., S.K.
Supervision: G.B.

