## Supplemental Figure 1 for "BCG vaccination of Diversity Outbred mice induces cross-reactive antibodies to SARS-CoV-2 spike protein"

### Supplemental Materials:

#### Creating a Standard Curve and Optimizing In-house ELISA:

Mouse mAb 240C has been widely documented as an antibody against the spike protein of the first SARS-CoV, and multiple studies demonstrate its ability to bind the RBD of the spike protein on SARS-CoV-2 as well [17]. We developed a standard curve using mAb 240C against plated rSpike protein in order to estimate the concentration of cross-reactive IgG in the serum of BCG-vaccinated mice. 240C standard curves ranged from a starting concentration of 10,000 pg/mL followed by a 2-fold dilution series, as the sera antibody concentrations of interest were located near the limit of detection. A limit of detection of 64 pg/mL was calculated for the 240C standard curve based on six standard curves (Supplemental Figure 1). Minimal variation in optical density across replicates was noted.

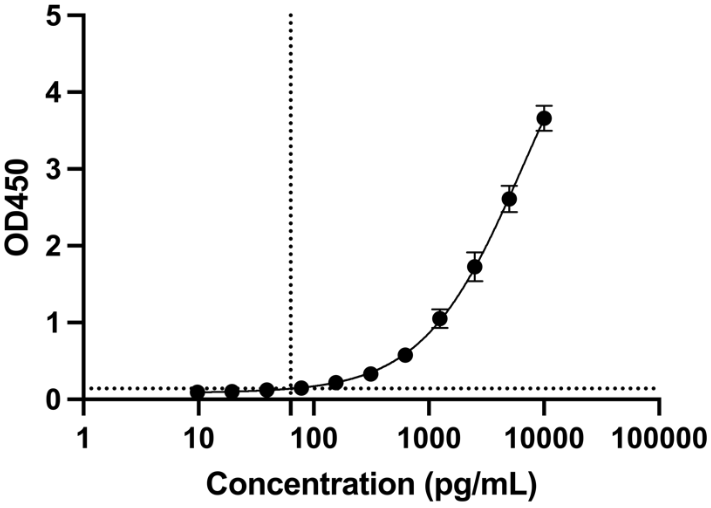


**Supplemental Figure 1:** Standard Curve using mAb 240C against rSpike. Standard curve was generated based on six replicates using aliquots of mouse monoclonal antibody 240C with a calculated starting concentration of 10,000 pg/mL. Error bars describe the variation in optical density across replicates. The limit of detection (dotted lines) was calculated to be 64 pg/mL.
